# Endophilin B1 primes mitochondria for execution

**DOI:** 10.64898/2026.06.25.733981

**Authors:** Shu-Chieh Susie Chang, Arni Thorlacius, Anna Sundborger-Lunna

## Abstract

Intrinsic apoptosis, or programmed cell death, is a vital response to stress and DNA damage in cells, and dysregulation of this pathway is common in cancers. The release of pro-apoptotic factors from mitochondria by the pro-apoptotic protein Bax is preceded by changes in the membrane properties of the outer mitochondrial membrane. We find that the membrane remodeling protein endophilin B1 primes membranes rich in the mitochondria-specific lipid cardiolipin for Bax-mediated membrane permeabilization, *via* a dual regulatory mechanism. We also show evidence that endophilin B1 translocates to the surface of mitochondria to co-localize with Bax during apoptosis *in situ*, where it forms biomolecular condensates.

## Introduction

Apoptotic outer mitochondrial membrane (OMM) permeabilization commits cells to enter the intrinsic apoptotic pathway by releasing intermembrane space proteins including cytochrome *c*, SMAC/DIABLO, and Omi/HtrA2 into the cytosol to activate caspases.^1–4^ Bax, alongside Bak, triggers OMM permeabilization through a tightly regulated multi-step process involving cytosolic-to-mitochondrial translocation,^5,6^ N-terminal conformational exposure,^7,8^ membrane insertion,^9,10^ oligomerization into higher-order homo-and hetero-complexes, and formation of discrete protein-lined membrane pores.^11,12^ This activation is governed by both protein regulators and the mitochondrial membrane milieu: BH3-only proteins abolish inhibition by pro-survival Bcl-2 family members and directly promote Bax/Bak oligomerization,^7,13,14^ while the lipid microenvironment, particularly cardiolipin (CL) oxidation and exposure on the OMM, critically dictates the kinetics and efficiency of Bax conformational activation, membrane integration, and pore assembly.^15–18^ Among mitochondrial lipids, CL has emerged as a critical determinant of Bax targeting and activation, supporting the idea that the membrane itself can function as a regulatory platform.^19–22^

Endophilin B1 (EnB1), also known as Bax-interacting factor 1 (Bif-1), is a Bin/Amphiphysin/Rvs167 (BAR) domain protein.^23^ EnB1 promotes membrane curvature and tubulation,^24^ consistent with its ability to shape lipid bilayers and maintain mitochondrial morphology.^25^ Initially identified as a Bax-interactor through yeast two-hybrid (Y2H) screening,^26,27^ EnB1 was subsequently found to delay Bax mitochondrial recruitment during apoptosis when knocked down.^28^ *In vitro* liposome permeabilization assays revealed that EnB1 promotes Bax-mediated membrane permeabilization (Bax-MMP) in a CL-, N-BAR domain-, and physical proximity-dependent manner.^29^ These findings place EnB1 at a strategically important position in apoptosis, where membrane remodeling and cell death signaling converge.

Our cryo-electron microscopy (Cryo-EM) structure of EnB1, bound to the surfaces of bicelles containing tetramyristoyl CL (14:0 CL), reveals that the N-terminal amphipathic helix 0 (H0) of EnB1 can be divided into two structurally distinct sections: A N-terminal portion that flexibly probes the surface of membranes, and a helical C-terminal portion that anchors the protein to target membranes.^30^ We also determined that EnB1 can destabilize lipid bilayers that contain tetraoleoyl CL (18:1) and suggested that the role of EnB1 during apoptosis is to destabilize the OMM following pro-apoptotic stimuli and cluster CL to promote Bax binding and OMM permeabilization.

Despite this progress, a central question remains unresolved: how does EnB1 modulate Bax-MMP at the mechanistic level? Specifically, it is not known whether EnB1 primarily promotes Bax by direct OMM recruitment or by altering the lipid organization and physical properties of the membrane. By defining how EnB1 shapes Bax activation at the membrane, this study aims to clarify a key mechanism linking mitochondrial membrane organization to apoptotic pore formation.

## Main

### Cardiolipin acyl chain saturation affects membrane phase

Liposomes containing tetrasteroyl CL (40% DOPC/35% DOPE/25% 18:0 CL/0.01% Laurdan) displayed significantly different generalized polarization (GP) of the membrane phase-sensitive dye Laurdan^31^ than CL-free liposomes (53% DOPC/47% DOPE /0.01% Laurdan; Figure 1a). Liposomes containing saturated CL had higher Laurdan GP, which suggests that the phase of the membrane was more rigid and gel-like than that of the membrane of vesicles containing no CL. This was also true for liposomes containing monounsaturated CL (40% DOPC/35% DOPE/25% 18:1 CL/0.01% Laurdan), however, the phase of 18:1 CL-liposomes was significantly more fluid and liquid-like than the phase of 18:0 CL-liposomes (Figure 1a). We observed no significant difference in Laurdan GP between CL-free liposomes and those that contained bovine heart CL (40% DOPC/35% DOPE/25% bovine heart CL), where the dominant form is tetralinoleoyl CL (18:2).

**Figure 1.**
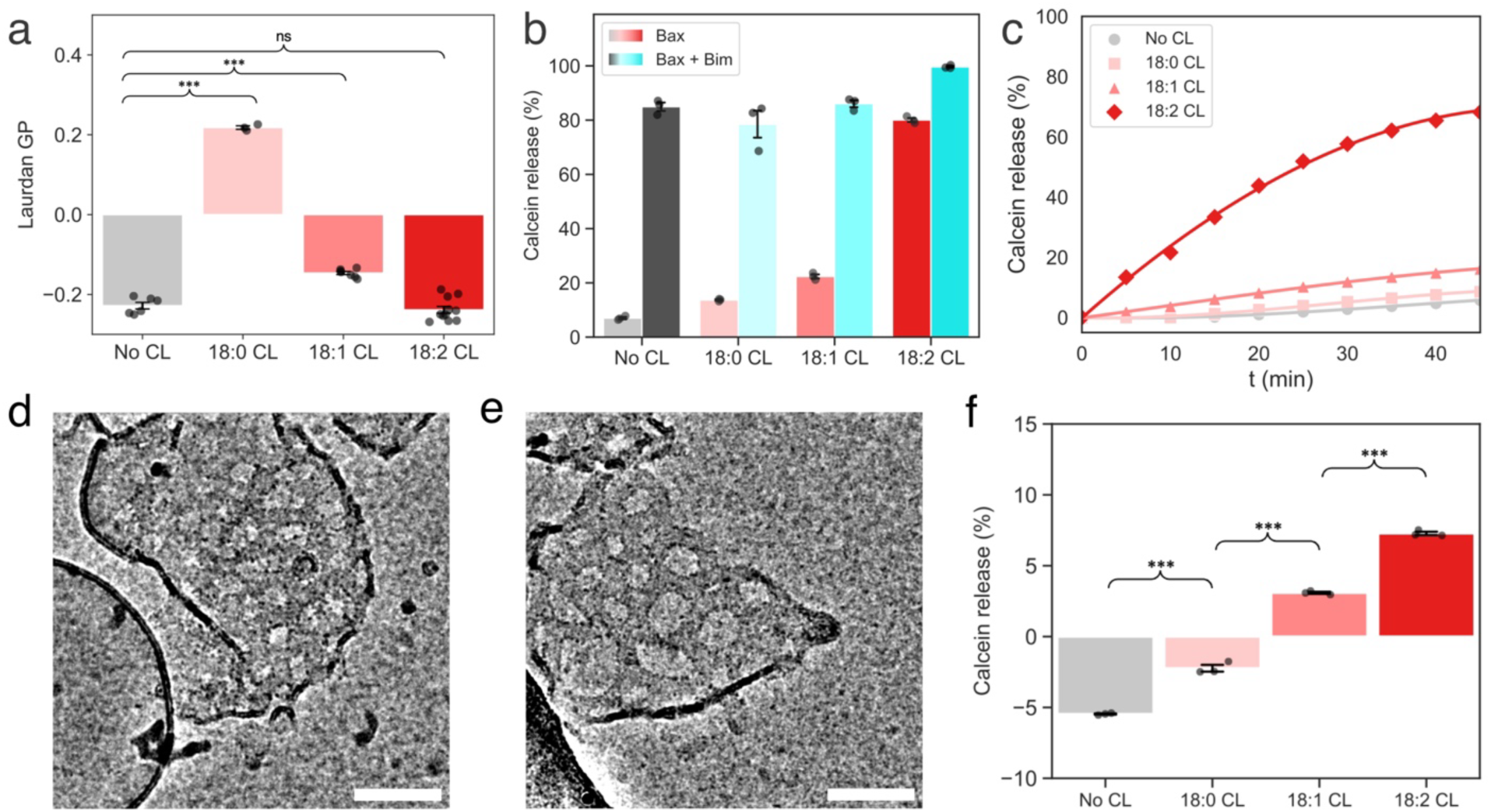
Bax and EnB1 disturb the bilayer of membranes containing unsaturated cardiolipin. (a) Laurdan GP for lipid compositions DOPC/DOPE (no CL) or supplemented with 25% CL (18:0, 18:1 or 18:2 CL) at 37 °C. Data represent mean ± SEM (n≥3). (b) Calcein release from liposomes composed of DOPC/DOPE (no CL) or supplemented with 25% CL (18:0, 18:1 or 18:2 CL) upon incubation for 30 minutes at RT with Bax alone or with Bim_BH3. Data represent mean ± standard deviation from the mean (SD; n = 3). (c) Time course of calcein release from liposomes containing no CL, 18:0 CL, 18:1 CL, or 18:2 CL upon Bax treatment. Cryo-EM image of 18:2 CL-liposomes following Bax incubation in the absence of Bim_BH3 (d) and in the presence of Bim_BH3 (e). Both reveal pore-like membrane structures. Scale bars represent 100 nm. (f) Calcein release after 30 minutes at room temperature from CL-free liposomes or CL-liposomes (18:0, 18:1 or 18:2 CL) upon incubation with EnB1 alone. Statistical significance is indicated by not significant (ns) if p > 0.05, *p < 0.05, **p < 0.01, or ***p < 0.001.

### Bax and EnB1 preferentially target polyunsaturated cardiolipin membranes

Given that CL is externalized to the OMM during apoptosis,^32,33^ and that EnB1 has been shown to preferentially bind CL,^30^ we investigated whether shared CL specificity provides a molecular basis for Bax-EnB1 co-localization. CL-free liposomes were compared to DOPC/DOPE liposomes supplemented with 25% heart CL (18:2 CL), a concentration previously shown to maximize assembly and GTPase activity of the mitochondrial fission protein Drp1, another CL-sensitive protein.^34^

BH3 motif-enhanced Bax-MMP of liposomes with encapsulated calcein was comparable between lipid compositions (Figure 1b). For CL-free liposomes, the membrane was <10% permeable to the fluorescent dye calcein, but when a BH3 activator was included (Bim BH3 motif peptide; Bim_BH3), significantly more encapsulated calcein was released. However, a majority of encapsulated calcein was released from 18:2 CL-liposomes, even in the absence of Bim_BH3. Cryo-EM revealed the presence of large membrane pores in CL-rich liposomes upon Bax incubation (Figure 1d). These were indistinguishable from pores observed in samples containing both Bax and Bim_BH3 (Figure 1e).

To assess whether this BH3-free activity depends on the acyl-chain saturation of CL, we measured Bax-MMP in the presence of CL-free liposomes or liposomes containing 25% 18:0, 18:1, or 18:2 CL (Figure 1b). Bax induced robust permeabilization of 18:2 CL-liposomes on its own, reaching ∼80% calcein release with faster kinetics than what was observed for saturated or monounsaturated CL-variants (Figure 1c). EnB1 displayed similar selectivity, through enhanced binding and permeabilization of 18:2 CL-liposomes (Figure 1f). Together, these results indicate that both Bax-MMP and EnB1 membrane binding preferentially occurs in the presence of polyunsaturated CL, consistent with a shared lipid-dependent basis for their co-localization at mitochondrial membranes.

### EnB1 increases membrane rigidity prior to and during tubulation

To determine whether EnB1 modulates Bax activity through membrane biophysical changes, we titrated EnB1 onto 18:2 CL-liposomes and monitored changes in Laurdan GP. At low EnB1 concentrations, Laurdan GP values increased significantly, indicating membrane rigidification (Figure 2a). We speculated that membrane rigidification was a result of EnB1 tubulation. Interestingly, we find that even sub-tubulation concentrations of EnB1 (sufficient to trigger permeabilization) induce membrane stiffening. Laurdan GP values reached a plateau when tubulation occurred (Figure 2d), suggesting that binding, and not necessarily helical scaffold assembly, of EnB1 is sufficient to promote changes in membrane phase behavior. When Bax was added to the same liposome sample we observed a similar change in Laurdan GP, i.e. there was no significant difference in the Laurdan GP of 1 μM EnB1 *versus* 1 μM of Bax (Supplementary Figure 1). This suggests that both EnB1 and Bax CL-membrane binding, at concentrations that trigger permeabilization, lead to membrane stiffening. When the two were added at the same time Laurdan GP significantly shifted compared to either one alone but there was no significant difference between both proteins added together *versus* 2 μM EnB1. This could mean that, while both proteins independently promote membrane rigidity, they do not act synergistically.

**Figure 2.**
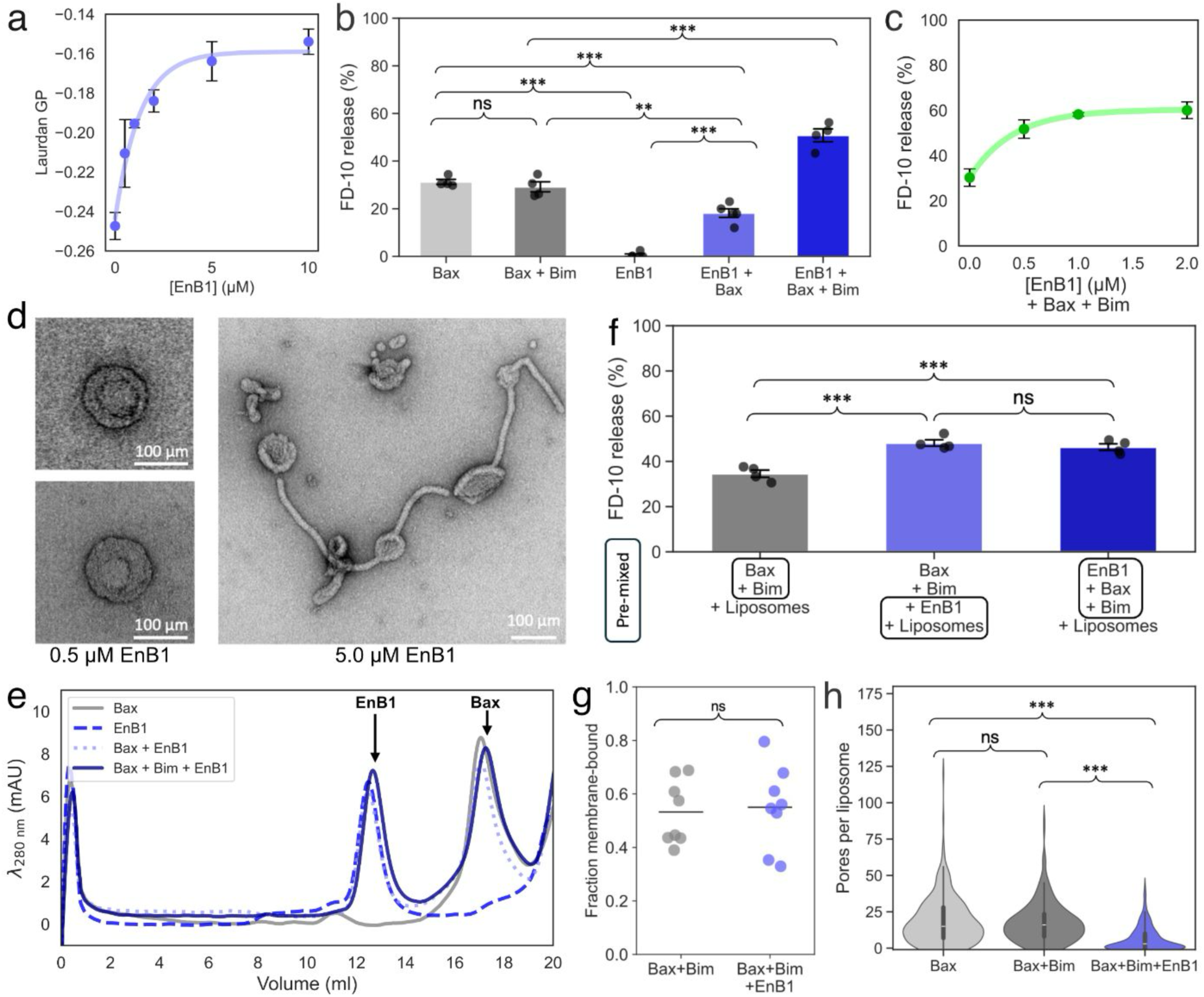
EnB1 differentially regulates Bax-MMP in a concentration-and context-dependent manner. (a) Laurdan GP increased as EnB1 was titrated onto 18:2 CL-liposomes (containing 0.5% Laurdan). Data represent mean ± standard error of the mean (SEM; n = 3). (b) FD-10 release from 18:2 CL-liposomes upon incubation with Bax, Bax+Bim_BH3, EnB1, EnB1+Bax, or EnB1+Bax+Bim_BH3. Data represent mean ± SD (n = 5). (c) EnB1 promoted BH3-enhanced Bax-MMP in a concentration-dependent manner. Liposomes were incubated with Bax+Bim_BH3 and increasing concentrations of EnB1 (0–2 μM). Data represent mean ± SD (n = 3). (d) Negative-stain EM images of 18:2 CL-liposomes incubated with EnB1 at two different concentrations. At 0.5 μM, liposomes remained predominantly spherical with EnB1 oligomerized side-to-side in a circle, whereas the 5 μM EnB1 sample was characterized by extensive membrane tubulation due to end-to-end polymerization and helical scaffold formation. Scale bars represent 100 nm. (e) Size-exclusion chromatography profiles of Bax, EnB1, and their mixtures in the absence or presence of Bim_BH3 using a Superdex 200 Increase 10/300 GL column. (f) FD-10 release from 18:2 CL-liposomes following sequential addition of proteins. 1^st^: Bax+Bim_BH3 were added together to liposomes. 2^nd^: Liposomes were incubated with EnB1 prior to the addition of Bax+Bim_BH3. 3^rd^: Bax+Bim_BH3+EnB1 were mixed before being added to liposomes. Data represent mean ± SD (n = 4). (g) There was no significant difference in the fraction of membrane-bound Bax between Bax+Bim_BH3 and Bax+Bim_BH3+EnB1. (h) The number of pores present per 18:2 CL-liposome (n) was compared between 150 cryo-EM micrographs each of Bax (n = 309), Bax+Bim_BH3 (n = 263) and Bax+Bim_BH3+EnB1 (n = 347). Statistical significance is indicated by ns if p > 0.05, *p < 0.05, **p < 0.01, or ***p < 0.001.

### EnB1 exhibits dual regulation of Bax-MMP on CL membranes

To examine the functional interplay between EnB1 and Bax on CL-rich membranes, we measured the release of 10 kDa FITC-dextran (FD-10), which is similar in size to cytochrome *c*, from 18:2 CL-liposomes in the presence of Bax and EnB1, with or without Bim_BH3. The use of FD-10 minimized the direct permeabilization contribution of EnB1, allowing us to focus on Bax-MMP. Bax induced substantial FD-10 release under these conditions on its own, consistent with its BH3-free ability to release calcein from CL-rich membranes (Figure 1b). In the absence of the Bim_BH3, EnB1 reduced Bax-MMP, indicating that EnB1 suppresses BH3-free Bax-MMP (Figure 1b). In contrast, when Bim_BH3 was present, EnB1 significantly enhanced Bax-MMP (Figure 2b; Supplementary Figure 2a).

To elucidate the mechanisms underlying this promotion effect, we performed EnB1 titration experiments. Increasing EnB1 concentrations progressively led to increased Bax-MMP of liposomes with encapsulated FD-10 in the presence of Bim_BH3, demonstrating a concentration-dependent stimulatory effect (Figure 2c). This enhancement reached a maximum at intermediate EnB1 concentrations (2 μM) and did not significantly increase at higher concentrations. Notably, the higher protein concentration range corresponded to conditions in which EnB1 induces membrane tubulation (at that lipid concentration), as shown by negative-stain EM (Figure 2d). Thus, the ability of EnB1 to facilitate BH3-enhanced Bax-MMP correlates directly with membrane rigidification, establishing that membrane biophysical conditioning by EnB1 is responsible for promotion of Bax-MMP, rather than EnB1 abundance or helical scaffold formation.

### EnB1 promotion of Bax-MMP happens at the membrane surface

To determine whether EnB1 stimulates Bax-MMP through direct protein–protein interaction or through changes in membrane properties, we examined whether these proteins form a stable complex in solution. Size-exclusion chromatography of EnB1, Bax+Bim_BH3 and EnB1+Bax+Bim_BH3 showed that each component eluted as a distinct species, indicating that the stimulatory effect of EnB1 does not require a stable soluble complex (Figure 2e). We then explored whether EnB1 can enhance Bax-MMP after a membrane-conditioning step. 18:2 CL-liposomes were pre-incubated with EnB1 before addition of Bax and Bim_BH3. Under these conditions EnB1 retained its ability to potentiate BH3-enhanced Bax-MMP (Figure 2f). This effect was comparable to when all components were mixed together before the addition of liposomes. This suggests that EnB1 promotion of Bax-MMP occurs at the membrane surface rather than in solution, and also that the way in which EnB1 potentiates Bax-MMP occurs rapidly and permanently upon EnB1 binding to the membrane.

### EnB1 enhances Bax pore-forming activity without affecting Bax membrane recruitment

We next investigated whether EnB1 promotes Bax-MMP by increasing Bax recruitment to liposomes. Sedimentation assays showed no statistical significance in the amount of membrane-bound Bax when either Bax+Bim_BH3 or Bax+Bim_BH3+EnB1 was added to 18:2 CL-liposomes under the same conditions where permeabilization was promoted (Figure 2g). We used cryo-EM to determine whether EnB1 produced changes in Bax-MMP. We collected cryo-EM micrographs at 100,000x magnification to visualize potential morphological or quantifiable differences. Large membrane pores were observed in all sample conditions (Supplementary Figure 3). Curiously, we found that there were significant differences in the number of pores per liposome between conditions. Despite causing the most efficient membrane permeabilization, Bax+Bim_BH3+EnB1 liposomes exhibited significantly fewer recognizable pores per liposome when compared to Bax or Bax+Bim_BH3 liposomes (Figure 2h). As expected, there was no significant difference in the number of pores per liposome when Bax and Bax+Bim_BH3 on 18:2 CL-liposomes were compared.

### Both H0 and BAR domain of EnB1 are required for effective promotion of Bax-MMP

Our previous work shows that H0 inserts into CL-liposomes to promote membrane destabilization and permeabilization.^30^ To define the structural determinants underlying EnB1-mediated enhancement of Bax activity, we analyzed several EnB1_H0 truncation and domain constructs. Deletion of the unstructured N-terminal segment of H0 (EnB1_ΔH0 1–10 aa/EnB1_ΔToe; Figure 3a) had little effect on the stimulatory activity of EnB1 (Figure 3b), which was surprising as a previous study concluded that this region was responsible for interactions with Bax in Y2H screens.^27^ In contrast, deletion of the alpha-helical segment of H0 (EnB1_ΔH0 11–31 aa/ EnB1_ΔHeel; Figure 3a) significantly decreased EnB1 promotion of BH3-enhanced Bax-MMP (Figure 3b), suggesting that this region is critical for the stimulatory effect. This pattern was mirrored in sedimentation assays (Figure 3c), to assess membrane binding ability, where there was a significant difference between the wild-type protein and H0 truncation mutants.

**Figure 3.**
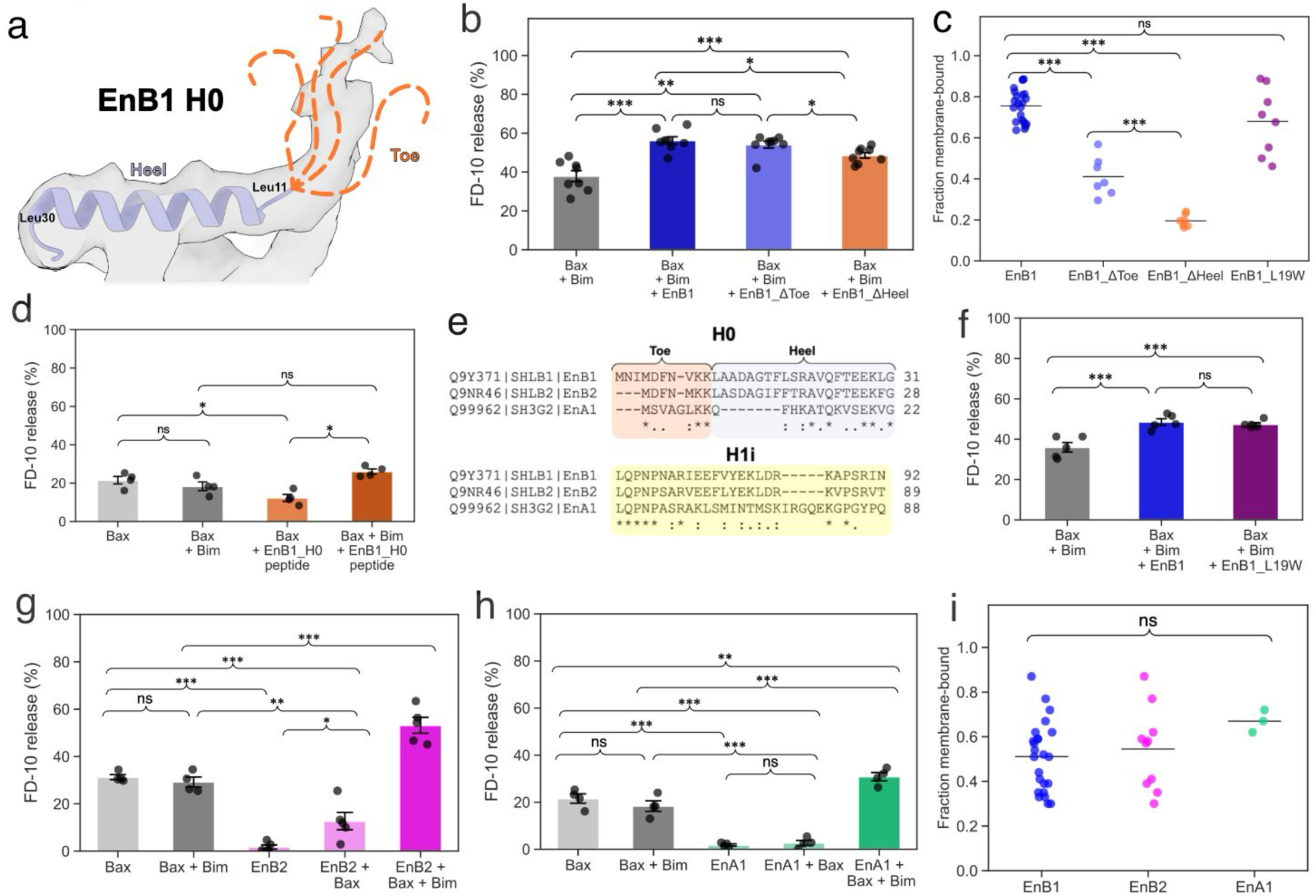
The H0 helix and BAR-domain scaffold cooperate to maximize EnB1 promotion of BH3-enhanced Bax-MMP. (a) In the cryo-EM structure of membrane-bound EnB1 (PDB: 9G2R; EMDB: 50981) the amphipathic H0 domain of EnB1 could be divided into two sections: a disordered N-terminus “toe” and an alpha-helical “heel”. (b) FD-10-encapsulated 18:2-CL liposomes were incubated with Bax+ Bim_BH3 in the absence or presence of EnB1, EnB1_ΔToe or EnB1_ΔHeel or EnB1_H0-heel. Data are shown as mean ± SEM (n = 8). (c) Deletions in H0 significantly impacted the membrane-binding ability of EnB1 mutants unlike the H0 mutation L19W. (d) Compared to EnB1, isolated EnB1_H0-heel was insufficient to fully reproduce the enhancement of Bax-MMP on 18:2 CL-liposomes. Data are shown as mean ± SEM (n = 5). (e) The L19W mutation had no significant effect on EnB1 promotion of Bax-MMP on 18:2 CL-liposomes. Data are shown as mean ± SEM (n = 5). **Endophilin-mediated Bax regulation is conserved across endophilin family members and independent of BH3-like motifs.** (f) Multiple sequence alignment of the amphipathic regions of endophilin family members EnB1, EnB2 and EnA1. (g) FD-10 release from 18:2 CL-liposomes upon incubation with Bax, Bax+Bim_BH3, EnB2, EnB2+Bax, or EnB2+Bax+Bim_BH3. Data represent mean ± SEM (n = 5). (h) FD-10 release from 18:2 CL-liposomes upon incubation with Bax, Bax+Bim_BH3, EnA1, EnA1+Bax, or EnA1+Bax+Bim_BH3. Data represent mean ± SEM (n = 5). (i) There was no significant difference in the binding of EnB1, EnB2 or EnA1 to 18:2 CL-liposomes in the presence of Bax+Bim_BH3. Data represent mean ± SEM (n = 5). Statistical significance is indicated by ns if p > 0.05, *p < 0.05, **p < 0.01, or ***p < 0.001.

Notably, isolated EnB1_H0-heel peptide (aa 11–31) significantly inhibited Bax-MMP when added to liposomes (EnB1_H0+Bax, Figure 3d), although the peptide was not sufficient to reproduce the full promotion effect when added together with the Bim_BH3 (EnB1_H0 +Bax+Bim_BH3, Figure 3d). Together, these results indicate that efficient promotion of BH3-enhanced Bax-MMP requires both the H0-mediated membrane binding and destabilization and the BAR-domain membrane scaffolding function.

### Endophilin-mediated Bax regulation is conserved across family members

Through sequence analysis, we identified two overlapping BH3-like motifs within EnB1 H0 (Supplementary Table 1), one forward and one in the reverse direction. *In silico* docking using CABS^35^ and AlphaFold3^36^ of EnB1_H0 into Bax resulted in models where H0 was predicted to bind the same BH3 binding groove as the Bid BH3 motif, as was reported in a 2013 study (Supplementary Figure 4a).^14^ CABS predictions showed EnB1_H0 docked in both directions (Supplementary Figure 4b-c) but scored the reverse BH3 orientation higher. AlphaFold3 on the other hand only docked the peptide in the reverse BH3 orientation (Supplementary Figure 4d). To test whether the role of EnB1 in Bax-MMP was BH3-mimicry, we designed a mutant EnB1_L19W that should be too bulky to fit into the BH3 binding groove of Bax. Other endophilin family members have phenylalanine instead of leucine in that position according to multiple sequence alignment (Figure 3e). There was no significant difference in the membrane binding activity of EnB1_L19W compared to the wild-type protein (Figure 3c) and, surprisingly, no significant difference in the promotion of Bax-MMP (Figure 3f).

We also examined Endophilin B2 (EnB2), which has high sequence identity with EnB1 but lacks critical BH3 motif amino acids. EnB2 also did not interact with Bax during Y2H screening.^27^ Remarkably, EnB2 mirrored the ability of EnB1 to promote Bax: it inhibited BH3-free Bax-MMP and promoted BH3-enhanced Bax-MMP on 18:2 CL-liposomes (Figure 3g). Furthermore, this was also true for Endophilin A1 (Figure 3h), which has not been reported to localize to OMMs. When different endophilins were incubated with Bax there was no significant difference between the amount of endophilin that was attached to the membrane (Figure 3i) and no difference in the level of promotion of Bax-MMP when normalized to Bax+Bim_BH3 activity (Supplementary Figure 2b). This family-wide conservation of function demonstrates that endophilin-mediated Bax regulation operates indirectly through shared membrane-remodeling mechanisms rather than directly through BH3 mimicry. The conserved H0/N-BAR dependence reinforces that membrane biophysical conditioning by amphipathic helices, not specific protein-protein interactions, drives pathway-specific Bax regulation.

### EnB1 and Bax co-localize during apoptosis

Previous studies established that stimulated emission depletion (STED) super-resolution microscopy can visualize Bax mitochondrial localization in apoptotic cells.^37,38^ We therefore applied STEDycon imaging at ∼40 nm resolution to examine EnB1 and Bax spatial organization in HeLa CCL-2 cells, which had been treated with 20 μM Embelin for 4 hours to induce intrinsic apoptosis (Figure 4a).^39–41^ Apoptotic cells were identified by distinct Bax-positive *puncta* (green). We observed striking co-localization of EnB1 and Bax signals as overlapping *puncta* forming dot, arc, and ring structures (Figure 4b). These observations indicate that EnB1 accumulates with Bax oligomeric assemblies at discrete OMM sites in apoptotic cells.

**Figure 4.**
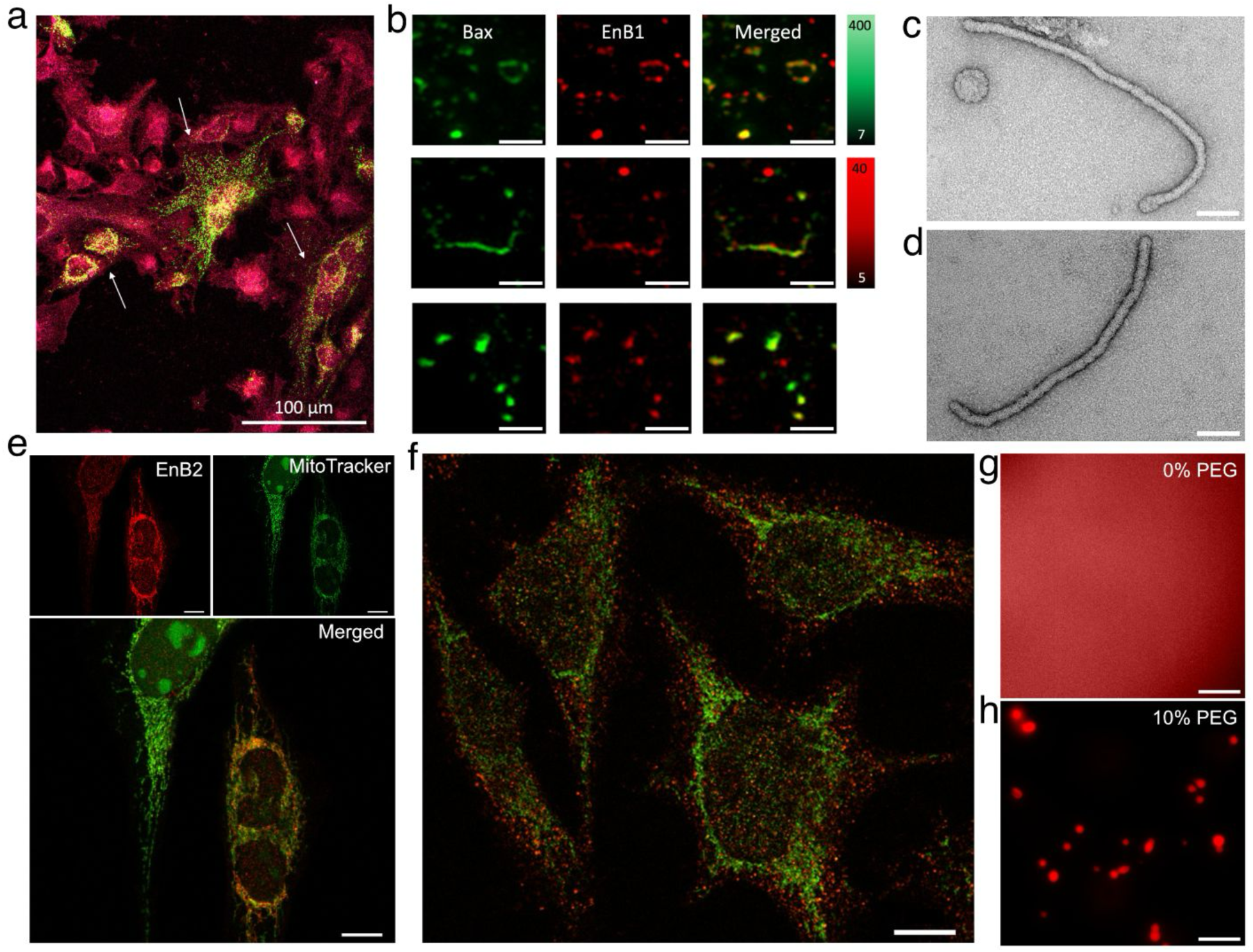
EnB1 co-localizes with Bax dot, arc, and ring structures during apoptosis. HeLa CCL-2 cells were treated with 20 μM Embelin for 4 h, immunostained for Bax (green) and EnB1 (red), and imaged by STED microscopy (∼40 nm resolution). (a) Overview showing apoptotic cells with Bax *puncta* (white arrows) versus non-apoptotic cells. The scale bar represents 100 μm. (b) High-magnification showing EnB1 colocalizing with Bax dot, arc, and ring structures. The scale bars represent 500 nm. (c) EnB1 tubulated liposomes are indistinguishable from (d) EnB2 tubulated liposomes. The scale bar represents 100 nm. **EnB2 is bound to the surface of mitochondria in healthy cells.** (e) Confocal images of healthy, untreated HeLa CCL-2 cells show the signal from immunolabeled EnB2 (red) and the signal from the mitochondrial marker MitoTracker™ (green) mostly overlap in the merged image. This indicates that EnB2 is constitutively bound to the surface of mitochondria. The scale bars represent 10 μm. (f) Confocal Image (STEDycon) of healthy, untreated HeLa CCL-2 cells immunostained for EnB1 (red) and EnB2 (green). The proteins localize to distinct places in the cell. **EnB1 forms phase-separated droplets *in vitro*.** Fluorescent signal for Atto-647N labeled EnB1 was diffuse at 18 μM concentration (g), however, addition of 10% PEG induced robust phase-separation of EnB1 at 18 μM (h). The scale bars represent 10 μm.

### EnB2 is present at the surface of mitochondria *in situ*

EnB2, like EnB1, readily tubulates liposomes *in vitro,* and EnB2 tubulated liposomes were indistinguishable from EnB1 tubulated liposomes when visualized using negative stain EM (Figure 4c-d). Confocal imaging (STEDycon) demonstrated that, in contrast to EnB1, EnB2 showed near-complete co-localization with the mitochondrial marker (MitoTracker™) in healthy, untreated HeLa CCL-2 cells (Figure 4e). This indicates that EnB2 constitutively localizes to the mitochondria. These findings were further supported by simultaneous dual-channel confocal imaging where we visualized untreated HeLa CCL-2 cells immunolabeled for EnB1 and EnB2 (Figure 4f), which demonstrated the differential subcellular distribution of the two proteins: EnB2 followed the elongated, reticular mitochondrial network, while EnB1 appeared as dispersed *puncta* throughout the cytoplasm with minimal overlap with the mitochondrial compartment. This differential localization suggests distinct functional roles for the two proteins, with EnB2 constitutively occupying the mitochondria and EnB1 being recruited from the cytoplasm to the OMM specifically during apoptosis. That would mean that endophilin B1 and B2 are distinctly regulated to control different aspects of mitochondrial function. Interestingly, EnB1 readily phase-separates in the presence of crowding agents (Figure 4g-h). Whether EnB1 *puncta* constitute condensates and whether these structures are involved in the regulation of EnB1 OMM remodeling, remains to be determined.

## Discussion

During early apoptosis, the changing composition and organization of the OMM takes center stage. Increased reactive oxygen species (ROS) production leads to the oxidation and externalization of CL to the OMM.^32,33^ Bax translocates to the OMM and permeabilizes it, leading to cytochrome *c* release and eventually cell death. Our results show that artificial membranes rich in CL become more fluid, which we interpret as an increase in packing defects, as the hydrophobic acyl chains of CL become more unsaturated (Figure 1a). Furthermore, we observe that CL acyl chain saturation directly influences Bax-MMP in the absence of BH3 activators (Figure 1b). We expect that membranes rich in oxidized CL will have even more packing defects and therefore be more vulnerable to Bax-MMP.

According to our observations, EnB1 is not present at high concentrations on healthy mitochondria *in situ* but translocates to the OMM and co-localizes with Bax, following treatment with Embelin (Figure 4a-b). In our *in vitro* experiments, EnB1 and Bax do not form stable complexes together in solution (Figure 2e) which agrees with previously published results,^42^ but both Bax and EnB1 preferentially bind membranes rich in polyunsaturated CL (Figure 1c and 1f). When we probed how the membrane phase was influenced by protein binding, we found that both proteins increased membrane rigidity (Supplementary Figure 1). Surprisingly, we find that EnB1 promotes Bax-MMP when added together with a BH3 activator but impedes it when no activator is present (Figure 2b). We observe less Bax-MMP when EnB1 is excluded despite there being no significant difference between the amount of membrane-bound Bax (Figure 2g). This suggests that EnB1 influences Bax pore formation rather than Bax membrane recruitment. Indeed, we observed fewer large pores during conditions where we expect EnB1 promotion of Bax-MMP in our cryo-EM data (Figure 2h). This could indicate that EnB1 modulates Bax distribution. Specifically, rather than Bax polymerizing into ever larger pores on a limited number of liposomes, the presence of EnB1 could promote the formation of smaller Bax pores across a broader population of liposomes (Figure 5). This widespread distribution would facilitate more efficient Bax-membrane binding, effectively “priming” the OMM for rapid Bax-MMP.

**Figure 5.**
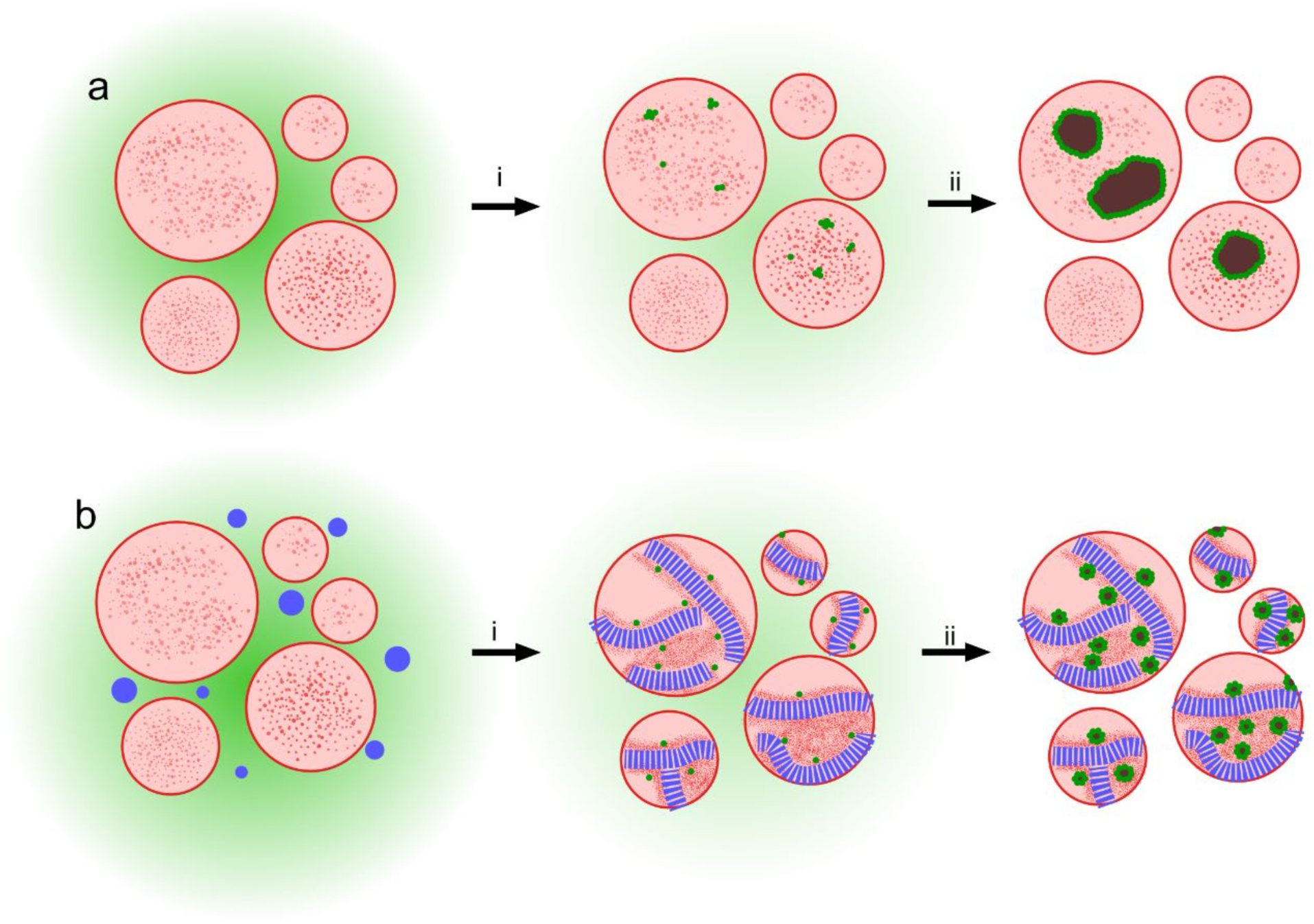
EnB1 promotion of Bax-mediated permeabilization. a) Stressed and fragmented mitochondria externalize oxidized CL. Here they are substituted for vesicles with surfaces rich in unsaturated CL (red dots). Bax (green) is in a diffuse and soluble state. i) Bax binds to the surface of the vesicles, inserts into the OMM recruiting more Bax from the cytosol to form oligomers. ii) Unchecked Bax polymerization results in large pores, the formation of which is favored due to Bax cooperative binding. Some vesicles are fully permeabilized while others are left intact. b) Vesicles surrounded by soluble Bax (green) and phase-separated droplets of EnB1. i) When CL becomes “available”, such as when oxidized CL is externalized by stressed mitochondria, EnB1 binds to the surface of the vesicles and oligomerizes. We propose that EnB1-mediated clustering of cardiolipin indirectly targets Bax to discrete areas on the OMM. ii) Steric hindrance due to EnB1 scaffolds would mean that the formation of smaller pores is favored, leading to more efficient Bax-targeting, pore formation and leakage from vesicles (and mitochondria).

The promotion we observed was independent of how long EnB1 had acted on the membrane (Figure 2f). This indicates that the mechanisms in which EnB1 primes the membrane for Bax-MMP happens rapidly and irreversibly. H0 mutant studies provided further evidence that EnB1 and Bax do not interact directly (Figure 3), however, the way in which H0 acts upon the membrane (when coordinated by the BAR domain) is critical for OMM permeabilization. We interpret this, together with the stiffening of the membrane, to be due to lipid partitioning driven by the insertion of amphipathic domains, which is supported by our previous model, wherein EnB1 plays a crucial role in regulating CL organization at the OMM by promoting membrane destabilization, and thereby, Bax-MMP.^30^

The fact that other endophilin family members, EnB2 and EnA1, can also promote Bax-MMP (Figure 3g and 3h) further reinforces the hypothesis that there is no specific interaction between EnB1 and Bax, and that the promotion effect has to do with membrane architecture and potentially Bax distribution. We unexpectedly observed that EnB2, not EnB1, is constitutively present at the surface of healthy mitochondria *in situ* (Figure 4e), which conflicts with several previously reported observations.^27,43–45^ EnB2 can purportedly form heterodimers with EnB1,^27^ and supposedly plays an important role during mitophagy.^44^ EnB2’s role at the surface of healthy mitochondria has not been studied.

We observe vesicle tubulation by EnB1 under conditions that lead to Bax-MMP, which highlights the close relationship between these states of membrane remodeling. We propose that at OMMs, a critical factor determining cell fate may be to coordinate membrane tubulation with Drp1-mediated fragmentation, a process that occurs during apoptosis. Tubulation could potentially also contribute to alternate downstream events, such as fragmentation of the OMM by EnB1 into vesicles or mitophagy related clearance of damaged mitochondria. Exactly what mechanisms dictate when and how EnB1 organizes on OMMs is unclear. We speculate that the local membrane lipid composition, i.e. the membrane architecture dictates the mode of EnB1-assembly, either into side-to-side or end-to-end scaffolds, which in turn controls how EnB1 will remodel the membrane. We predict that membranes rich in CL will favor negative curvature formation and resist tubulation, which may contribute to sequestration of CL by EnB1.

EnB1 phase separates *in vitro* (Figure 4g and 4h) and forms *puncta* consistent with condensate formation in healthy, untreated HeLa CCL-2 cells (Figure 4f). Results from a 2009 study indicate that Bax activates EnB1 oligomerization *in vitro* (after longer incubations at higher concentrations of the proteins than shown here),^42^ which could be related to EnB1 *in situ* condensate formation. It has been shown that EnA1 also forms condensates,^46^ which directly influences its ability to remodel the plasma membrane during synaptic vesicle recycling,^47^ and that EnA1 condensate formation is regulated by Ca^2+^ signaling.^48,49^ Intracellular Ca^2+^ signaling plays an important role in controlling organelle behavior. A 2014 study found mitochondrial Ca^2+^ influx to be a key driver of oxidative stress and ROS production during apoptosis.^50^ We speculate that EnB1 membrane remodeling activity is similarly regulated through phase separation and *puncta* formation, and that downstream membrane remodeling events involved in apoptosis and mitophagy are triggered by sudden increases in intracellular Ca^2+^ concentration.

## Materials and methods

### Bax expression and purification

Bax-intein-CBD (IMPACT™ system; New England Biolabs) was expressed recombinantly in *Escherichia coli* C41 (DE3) cells. Bacteria were grown in Terrific Broth (TB) and induced with 0.6 mM IPTG when the culture reached OD_600_ = 0.8, followed by expression at 20 °C for 16 hours. Cells were harvested and resuspended in Lysis buffer (20 mM HEPES, pH 7.4, 500 mM NaCl) supplemented with cOmplete™ Protease Inhibitor Cocktail (Sigma-Aldrich) and 10 μg/ml Bovine pancreas DNase I (Merck). Cells were lysed at 35 kPSI using a cell disruptor (Constant Systems), and the lysate was centrifuged at 30,000 *× g* for 60 min. Clarified lysate was applied to a gravity column containing chitin resin (New England Biolabs) pre-equilibrated with Lysis buffer. Intein self-cleavage was triggered through the addition of Lysis buffer supplemented with fresh 50 mM dithiothreitol (DTT; Merck) and incubation at 4 °C for 64 hours. Fractions were collected from the affinity resin and applied directly to a HiLoad® 16/600 Superdex® 75 pg column (Cytiva) pre-equilibrated with Bax sizing buffer (20 mM HEPES pH 7.4, 150 mM NaCl, 0.5 mM EDTA). Protein purity was estimated using SDS-PAGE and WB using the anti-Bax (2D2; mouse monoclonal IgG_1_ κ; Santa Cruz Biotechnology). Fractions containing monomeric Bax were stored at 4 °C and used within 2 weeks of purification.

### EnA1 and EnB2 cloning

**EnA1**. The *EnA1* coding sequence was amplified from iPSC-derived neuron cDNA by PCR using primers 5′-TCCTGCACCATGTCGGTG-3′ (forward) and 5′-GCCAGCCAGCATAACATC-CTAAT-3′ (reverse). The purified amplicon was subsequently subjected to a two-step PCR to append vector-homologous overhang sequences for Gibson assembly using primers 5′-CAGGGGCCCCATATGTCGGTGGCCGGCCTCAAGAA-3′ (forward) and 5′-GTGGTGGTGCTC-GAGCTAATGGGGCAGGGCAACCA-3′ (reverse). In parallel, the pET28a backbone incorporating a PreScission protease recognition site was linearized by inverse PCR using primers 5′-CATATGGGGCCCCTGGAACA-3′ (forward) and 5′-CTCGAGCACCACCACCACCA-3′ (reverse). Both PCR products were gel-purified and assembled using NEBuilder® HiFi DNA Assembly Master Mix (New England Biolabs) according to the manufacturer’s instructions. The assembled product was transformed into chemically competent *E. coli* Top10 cells, and correct insertion was confirmed by whole-plasmid sequencing (Eurofins Genomics).

**EnB2**. The *EnB2* coding sequence was amplified from HCT116 cell-derived cDNA by PCR using primers 5′-ACGCCATGGACTTCAACATG-3′ (forward) and 5′-GCTTGAGCTAGCTGAGC-AGT-3′ (reverse). The purified amplicon was subsequently subjected to a two-step PCR to append restriction site-compatible sequences using primers 5′-CAGGGGCCCCATATGTCG-GTGGCCGGCCTCAAGAA-3′ (forward) and 5′-GTGGTGGTGCTCGAGCTAATGGGGCAGGGC-AACCA-3′ (reverse). The resulting PCR product and the pET28a vector were each digested with NdeI and XhoI (FastDigest, Thermo Fisher Scientific), gel-purified, and ligated using T4 DNA Ligase (Thermo Fisher Scientific) according to the manufacturer’s instructions. The ligation product was transformed into chemically competent *E. coli* Top10 cells, and correct insertion was confirmed by whole-plasmid sequencing (Eurofins Genomics).

### Expression and purification of endophilins

His_12_-SUMO-endophilin-B1 (full-length and truncated forms), His_6_-endophilin-B2 and His_6_-endophilin-A1 were expressed recombinantly in *E. coli* Rosetta2 cells. Bacteria were grown in lysogeny broth (LB) and induced with 0.2 mM IPTG at OD_600_ = 0.6, followed by expression at 18 °C for 18 hours. Cells were harvested and resuspended in cell lysis buffer (20 mM Tris, 500 mM NaCl, 20 mM Imidazole, 0.5 mM TCEP, and 0.25% Triton X-100, pH 8.2) supplemented with cOmplete™ Protease Inhibitor Cocktail (Sigma-Aldrich) and 10 μg/ml Bovine pancreas DNase I (Merck). Cells were lysed at 35 kPSI using a cell disruptor (Constant Systems), and the lysate was centrifuged at 30,000 *× g* for 60 min. Clarified lysate was applied to a gravity column containing Protino® Ni-NTA agarose beads (MACHEREY-NAGEL) pre-equilibrated with Binding buffer (20 mM Tris, 500 mM NaCl, 20 mM Imidazole, 0.5 mM TCEP, pH 8.2). The column was washed with wash buffer (20 mM Tris, 500 mM NaCl, 50 mM Imidazole, 0.5 mM TCEP, pH 8.2) and endophilins were eluted with elution buffer (20 mM Tris, 500 mM NaCl, 500 mM Imidazole, 0.5 mM TCEP, pH 8.2). Eluted protein was dialyzed into Cleavage buffer (20 mM Tris, 300 mM NaCl, 0.5 mM TCEP, pH 8.0). For His-SUMO-EnB1 SUMO (Ulp1) protease was added to dialyzed protein at 1:10 (w/w) for overnight cleavage at 4 °C with gentle agitation. His_6_-tagged SUMO protease and uncleaved fusion protein were removed by applying the filtered sample to a Nickel affinity column and collecting the unbound fraction. For His_6_-endophilin-B2 and His_6_-endophilin-A1 the same was done except the tag was removed using His-GST-HRV 3C protease (PreScission Protease). Cleaved protein was concentrated and passed over a HiLoad® 16/600 Superdex® 200 pg column (Cytiva) using EnB1 sizing buffer (20 mM HEPES, 150 mM NaCl, 0.5 mM TCEP, pH 8.0). SDS-PAGE analysis was used to estimate the purity of the sample. Fractions containing pure endophilin (≥99% w/w) were concentrated to 2 mg/ml, flash frozen in liquid N2 and stored at−70 °C until use.

### Lipid film preparation

Lipid stocks dissolved in chloroform (Avanti Research), 18:1 DOPC (1,2-dioleoyl-sn-glycero-3-phosphocholine), 18:1 DOPE (1,2-dioleoyl-sn-glycero-3-phosphoethanolamine), 18:0 cardio-lipin (1’,3’-bis[1,2-distearoyl-sn-glycero-3-phospho]-glycerol), 18:1 cardiolipin (1’,3’-bis[1,2-dioleoyl-sn-glycero-3-phospho]-glycerol), or bovine heart cardiolipin (∼95% 18:2 CL; 1’,3’-bis[1,2-dilinoleoyl-sn-glycero-3-phospho]-glycerol) were mixed and dried under N_2_ gas to form dry lipid films. These were stored under vacuum overnight to ensure full removal of the solvent.

### Laurdan generalized polarization

Lipid films spiked with 0.5 mol% Laurdan (Avanti Research) were resuspended in Bax sizing buffer so that the concentration of Laurdan was at least 10 μM and extruded through a 0.4 µm filter at least 21 times. Laurdan liposomes were added to samples in a 96 well plate and the spectra of Laurdan recorded using a CLARIOstar Plus plate reader (BMG Labtech) with the excitation frequency set to 348 ± 10 nm measuring the emission intensity at 10 nm intervals between 400 ± 10 nm and 560 ± 10 nm. Laurdan generalized polarization (GP) was calculated using the following equation:

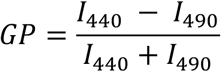

Where *I_440_* and *I_490_* represent the fluorescent intensity measured at 440 nm and 490 nm, respectively.

### Calcein permeabilization assay

Lipid films were solubilized in a 100 mM calcein (Merck) solution and extruded through a 0.4 µm filter at least 21 times. Calcein-encapsulating liposomes were separated from free calcein by gel filtration using G-50 Sephadex® resin (Cytiva) and Bax sizing buffer. The encapsulation efficiency for each batch was determined by comparing liposomes alone (negative control) with liposomes in the presence of 1% Triton X-100 (Alfa Aesar), which fully permeabilizes liposomes. The fluorescence signal from positive controls was at least 10-fold higher than that of negative controls in all subsequent assays.

Calcein-encapsulated liposomes were added to protein samples in a 96-well plate and increase in calcein fluorescence was monitored in a CLARIOstar Plus plate reader at excitation wavelength 482 nm and bandwidth 16 nm, emission wavelength 530 nm and bandwidth 40 nm with the dichroic mirror set to 504 nm. Sample fluorescence was either measured every 5 minutes up to 60 minutes or only once after a 20-minute incubation at room temperature. Every run had wells with calcein-encapsulated liposomes without protein (negative controls) as well as calcein-encapsulated liposomes supplemented with 1% Triton X-100, which represented the maximum release of calcein (positive controls).

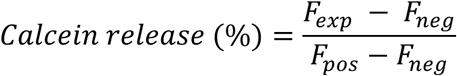

Where *F_exp_* is the fluorescence of the sample, *F_neg_* the average fluorescence of the negative controls and *F_pos_* the average signal from positive controls.

### FITC-dextran permeabilization assay

Lipid films were solubilized in a 10 mM Fluorescein isothiocyanate (FITC)–dextran (average MW 10,000; FD-10; Merck) solution and extruded through a 0.4 µm filter at least 21 times. FD-10 encapsulating liposomes were separated from free FITC-dextran by gel filtration using a Superose® 6 Increase 10/300 GL column (Cytiva) and Bax sizing buffer. The encapsulation efficiency for each batch was determined by comparing liposomes alone (negative control) with liposomes in the presence of 1% Triton X-100 (Alfa Aesar), which fully permeabilizes liposomes. The fluorescence signal from positive controls was at least 10-fold higher than that of negative controls in all subsequent assays.

FD-10-encapsulated liposomes were added to protein samples in 200 μl ultracentrifuge tubes. Samples were immediately centrifuged at 100,000 × *g* for 20 minutes in an Optima MAX-TL ultracentrifuge (Beckman Coulter) equipped with a TLA-100 fixed-angle rotor (Beckman Coulter). Release of FD-10 was compared between samples and controls by collecting the supernatant from each ultracentrifuge tube and transferring them separate wells in a 96 well plate and measuring FITC fluorescence intensity in a CLARIOstar Plus plate reader at excitation wavelength 482 nm and bandwidth 16 nm, emission wavelength 530 nm and bandwidth 40 nm with the dichroic mirror set to 504 nm. Every run had wells with FD-10-encapsulated liposomes without protein (negative controls) as well as FD-10-encapsulated liposomes supplemented with 1% Triton X-100, which represented the maximum release of FITC-dextran (positive controls).

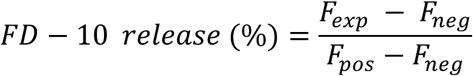

Where *F_exp_* is the fluorescence of the sample, *F_neg_* the average fluorescence of the negative controls and *F_pos_* the average signal from positive controls.

### Sedimentation assay

Lipid films were solubilized in Bax or EnB1 sizing buffers and extruded through a 0.4 µm filter at least 21 times. Liposomes were incubated with protein samples for at least 20 minutes before being pelleted *via* centrifugation at 100,000 *× g* for 20 minutes in an Optima MAX-TL ultra-centrifuge. The supernatant was removed and the pellet fraction re-suspended in fresh buffer. The re-suspended pellet fractions were used to run SDS-PAGE, together with samples taken prior to ultracentrifugation. The membrane-bound protein fraction was estimated by dividing the amount of protein present in the resuspended pellet with the amount of protein present in the sample prior to ultracentrifugation, using ImageJ^51^.

### Pore quantification assay

Lipid films were solubilized in Bax or EnB1 sizing buffers and extruded through a 0.2 µm filter 21 times. Liposomes were incubated with protein samples: 2.2 μM Bax + 400 μM 40% DOPC/35% DOPE/25% heart CL liposomes (1:185; Bax:lipid), 2.1 μM Bax + 0.5 μM Bim_BH3 + 400 μM 40% DOPC/35% DOPE/25% heart CL liposomes (4:1:800; Bax:Bim_BH3:lipid) and 0.5 μM EnB1 + 2.1 μM Bax + 0.5 μM Bim_BH3 + 400 μM 40% DOPC/35% DOPE/25% heart CL liposomes (1:4:1:800; EnB1:Bax:Bim_BH3:lipid). Samples were incubated at room temperature for at least 10 minutes before being applied to glow-discharged QUANTIFOIL® R 2/1 Holey Carbon grids (Quantifoil Micro Tools GmbH), blotted using filter paper (Whatman®) and plunge frozen in liquid ethane using a Vitribot Mark IV (Thermo Fisher Scientific). Cryo-EM movies were recorded at the Cryo-EM Uppsala facility using a 200 kV Glacios (Thermo Fisher Scientific) electron microscope equipped with a Falcon 4i direct electron detector (Thermo Fisher Scientific) and a Selectris Energy Filter (Thermo Fisher Scientific) where the slit diameter was set to 10 eV. Movies were imported, patch motion corrected, patch CTF corrected and manually curated in CryoSPARC^52^ to remove micrographs that did not contain liposomes. A script was generated to low pass filter the micrographs to 15 Å resolution (code available upon request). Pore quantification was performed in ImageJ by marking each pore with colored markers, one color per visible liposome, and then quantifying the number of dots per micrograph according to color.

### Cell culture

HeLa CCL-2 cells were maintained in Dulbecco’s Modified Eagle Medium (DMEM) supplemented with 10% fetal bovine serum (FBS), 100 U/mL penicillin, and 100 µg/mL streptomycin at 37°C in a humidified atmosphere containing 5% CO₂. Cells were sub-cultured by trypsinization at a 1:10 split ratio upon reaching ∼80% confluence and were discarded after the 20^th^ passage.

### STED super-resolution microscopy

Coverslips were cleaned by immersion in 99% ethanol for 1 hour, transferred to 6-well plates, and sterilized under UV light in a biosafety cabinet for 30 minutes prior to use. Cells were seeded at 5 × 10⁵ cells per well and cultured overnight to approximately 70% confluence. To label mitochondria, cells were incubated with pre-warmed (37°C) MitoTracker™ Orange CMTMRos (100 nM; Thermo Fisher Scientific) in growth medium for 30 minutes at 37°C. To induce apoptosis, cells were treated with 20 µM Embelin for 4 hours, followed by three washes with PBS. Cells were fixed with 4% paraformaldehyde for 10 minutes, rinsed three times with PBS, and permeabilized with 0.05% digitonin in PBS for 10 minutes. Non-specific binding was reduced by treatment with Image-iT® FX Signal Enhancer (Thermo Fisher Scientific) for 30 minutes, followed by blocking in 3% bovine serum albumin (BSA) in PBS containing 0.01% digitonin for 1 hour. Coverslips were then incubated overnight at 4°C with primary antibodies diluted in PBS containing 1% BSA and 0.01% digitonin: rabbit polyclonal anti-Bax (1:500; Cell Signaling Technology) and mouse anti-EnB1 (1:200; Santa Cruz Biotechnology). After three washes with PBS containing 0.01% digitonin (3 minutes each), coverslips were incubated with STAR RED-or STAR Orange-conjugated secondary antibodies (1:200; Abberior) for 1 hour at room temperature, followed by three additional washes as described. Staining quality was verified by widefield fluorescence microscopy prior to STED imaging.

STED images were acquired on a STEDycon (Abberior Instruments) mounted on an Olympus inverted microscope with a 100×/1.45 NA oil immersion objective. Excitation was performed using 561 nm and 640 nm pulsed diode lasers, with a 775 nm pulsed depletion laser. The pinhole was set to 64 µm, and images were acquired at a pixel size of 20 nm with a pixel dwell time of 10 µs and 25 line accumulations.

### Fluorescence microscopy

Wild-type EnB1 was covalently labeled with Atto-647N fluorescent dye (ATTO-TEC GmbH) using maleimide coupling. Purified protein was buffer exchanged to 1xPBS buffer (Merck) and incubated with dye at a molar ratio of 1:1 (dye:protein) for 24 hours at 4 °C. Labeled protein was separated from free dye by gel filtration using a Superdex® 200 Increase 10/300 GL column (Cytiva). Labeled protein was imaged at 18 μM, with and without 10% (w/v) PEG 8000 (Fluka Analytical), using a Zeiss Axio Observer epifluorescence microscope equipped with a Lumencor Spectra-X LED light source, a 100×, 1.4 NA oil immersion objective and a Andor Zyla sCMOS camera. Fluorescence images were acquired with an exposure time of 25 ms and nominal LED light source power of ∼5%.

## Acknowledgements

Data was collected at the Cryo-EM Uppsala facility at Uppsala university. Cryo-EM data processing was enabled by the Davinci cluster at Uppsala university ICM with generous technical assistance from Prof. Filipe Maia and Dr. Daniel Larsson. The BioVis platform of Uppsala University was used to conduct experiments using negative stain EM, supported by Monika Hodik, PhD and Karin Staxäng, TMA, as well as confocal and STED microscopy. Assoc. Prof. Tanel Punga of the Department of Medical Biochemistry and Microbiology; Infection and Immunity generously provided HeLa CCL-2 cells. We would also like to thank Ana Maria Villamil Giraldo, PhD and Steinar Mannsverk, PhD for their help with fluorescence microscopy.

## Supplementary Figures

**Supplementary Figure 1.**
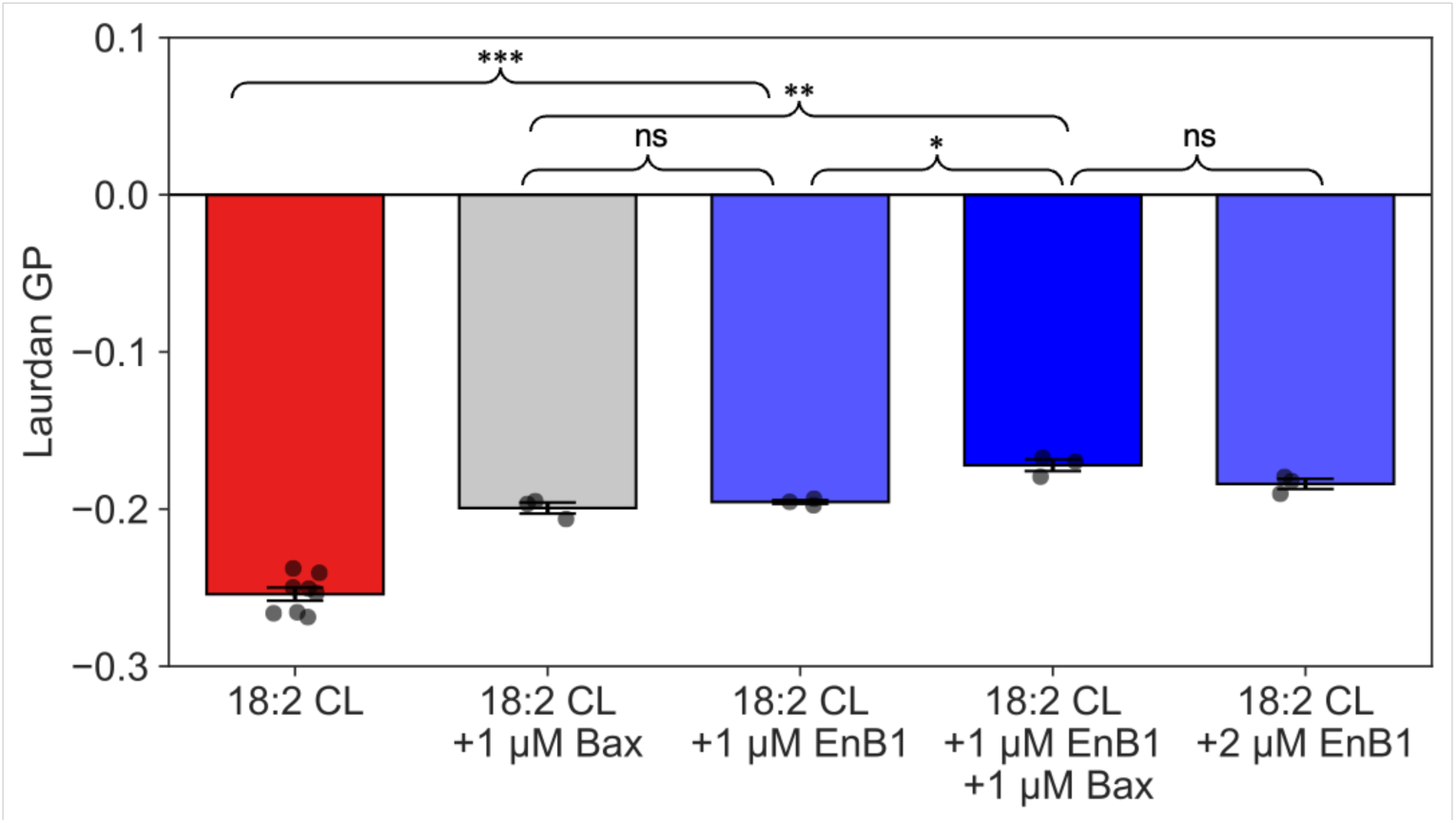
The addition of Bax and EnB1 to 18:2 CL-liposomes both increase laurdan GP, as well as when they are added simultaneously. Data represent mean ± SEM (n ≥ 3). Statistical significance is indicated by not significant (ns) if p > 0.05, *p < 0.05, **p < 0.01, or ***p < 0.001.

**Supplementary Figure 2.**
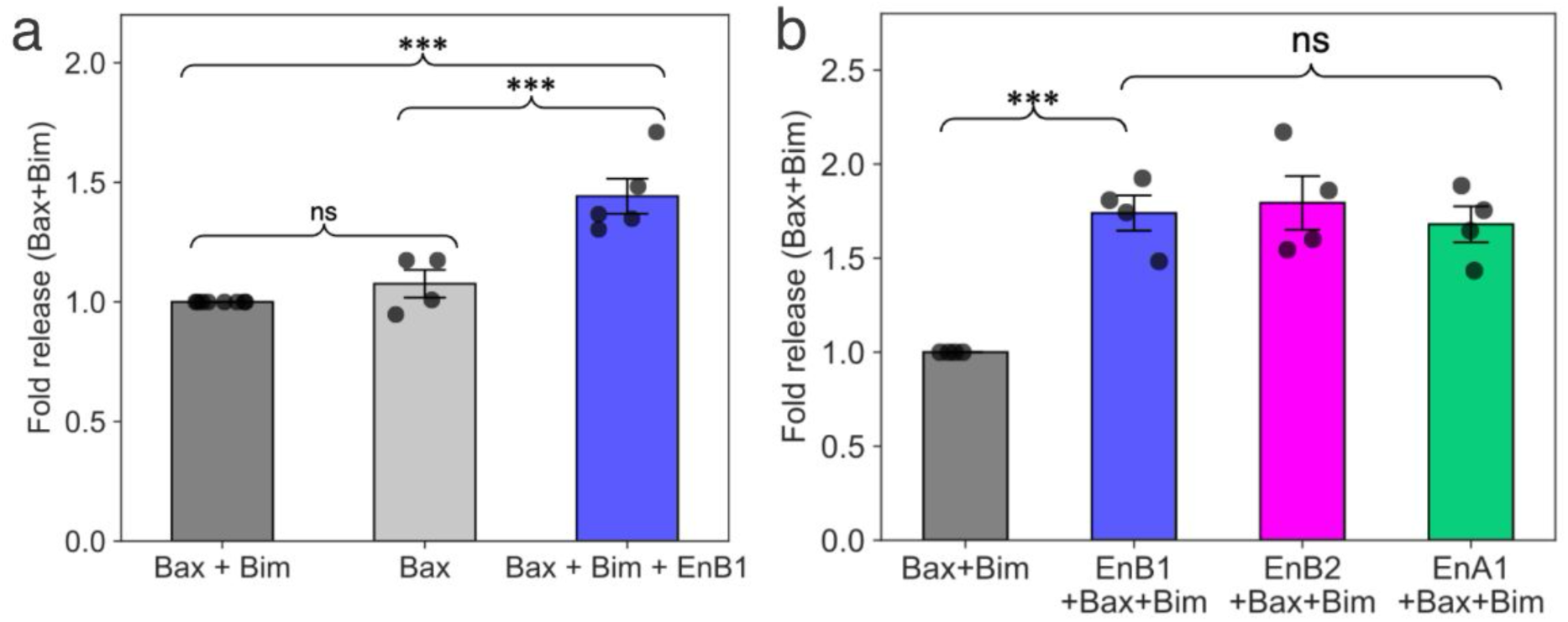
Validation of endophilin promotion of BH3-enhanced Bax-MPP. (a) Fold changes in FD-10 release due to Bax-MMP from 18:2 CL-liposomes were calculated relative to the condition containing Bax and Bim (Bax+Bim). The plot shows biological replicates, where each dot represents the average of at least three technical replicates. Bax alone showed no significant difference from Bax+Bim, while the condition containing Bax, Bim, and EnB1(Bax+Bim+EnB1) released significantly more FD-10 than Bax+Bim. Data represent mean of the biological replicates ± SEM (n ≥ 4) (b) Fold changes in Bax-mediated FD-10 release from 40% DOPC/35% DOPE/25% heart CL liposomes by the addition of different endophilins were calculated relative to the condition containing Bax and Bim (Bax+Bim). While there are significant differences in the fold FD-10 release compared to Bax+Bim for each individual endophilin there are no significant differences when the different endophilin samples are compared to each other. Data represent mean ± SEM (n = 4). Statistical significance is indicated by ns if p > 0.05, *p < 0.05, **p < 0.01, or ***p < 0.001.

**Supplementary Figure 3.**
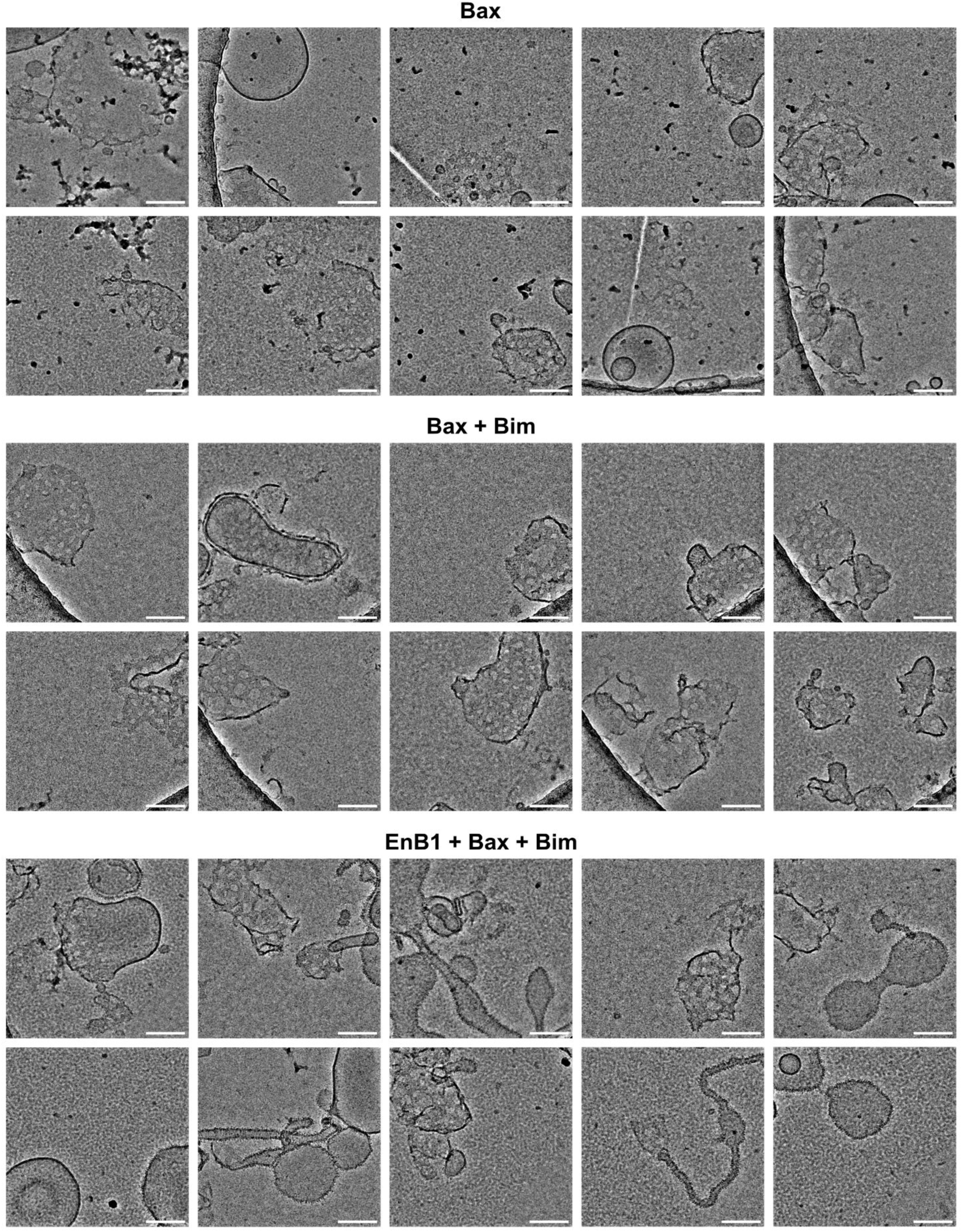
Examples of cryo-EM micrographs of 25% heart CL liposomes incubated with Bax, Bax + Bim or EnB1 + Bax + Bim which were used in the pore quantification assay whose results are shown in Figure 2h. The scale bars represent 100 nm.

**Supplementary Figure 4.**
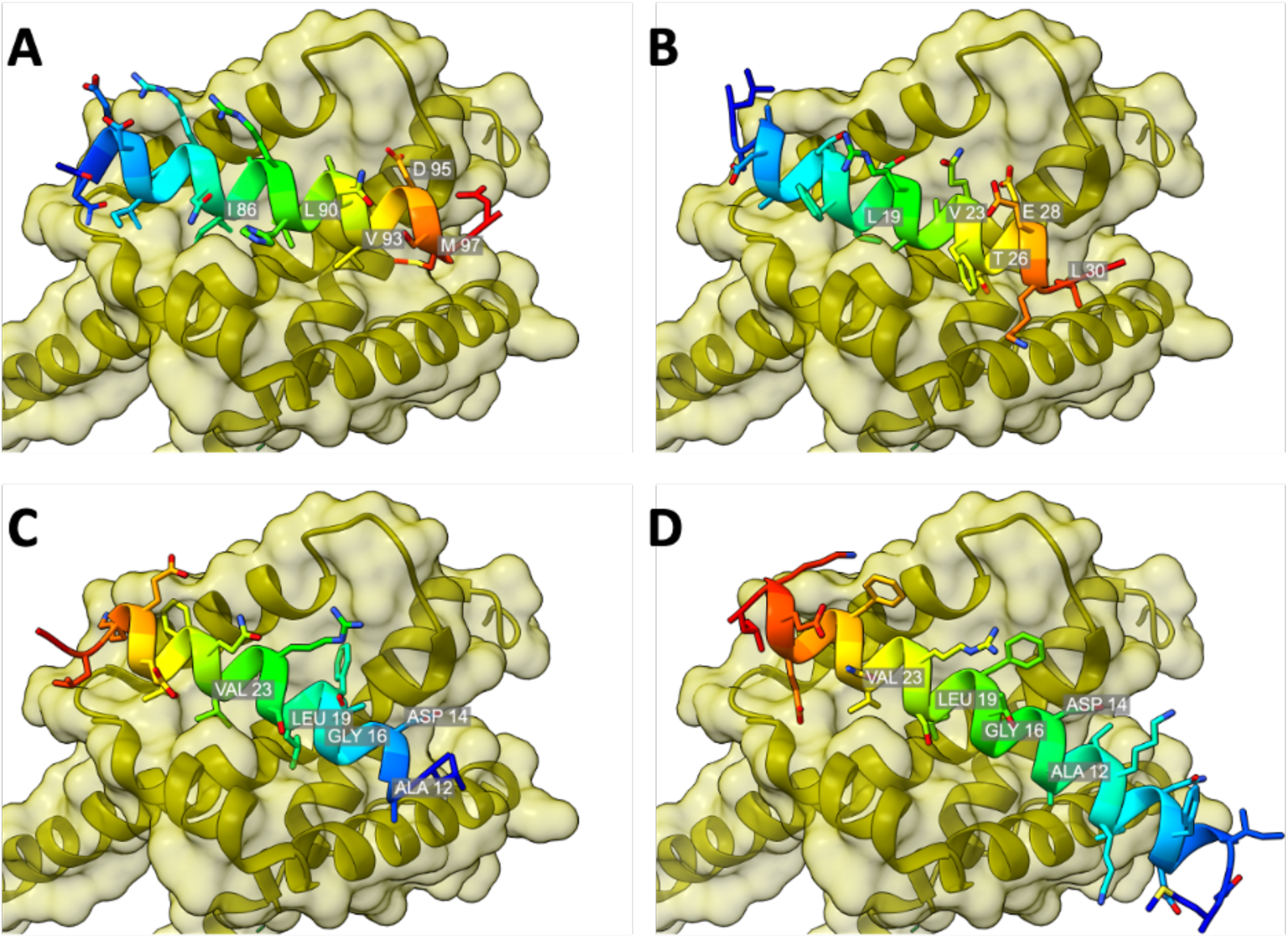
A) MiniBID BH3 peptide bound to the hydrophobic binding cleft of Bax (PDB ID: 4ZIG). CABS docking results show two possible binding orientations for a truncated H0 peptide (L11-G30): same orientation as the BID BH3 and other canonical BH3 motifs (B) or in the reversed orientation (C). D) The AlphaFold 3 prediction for full length H0 depicts the peptide bound to the hydrophobic groove of Bax in the reverse orientation. The labelled residues in all figures correspond to conserved BH3 motif residues that are shown in table 11 in bold.

## Supplementary Table

**Supplementary Table 1.**
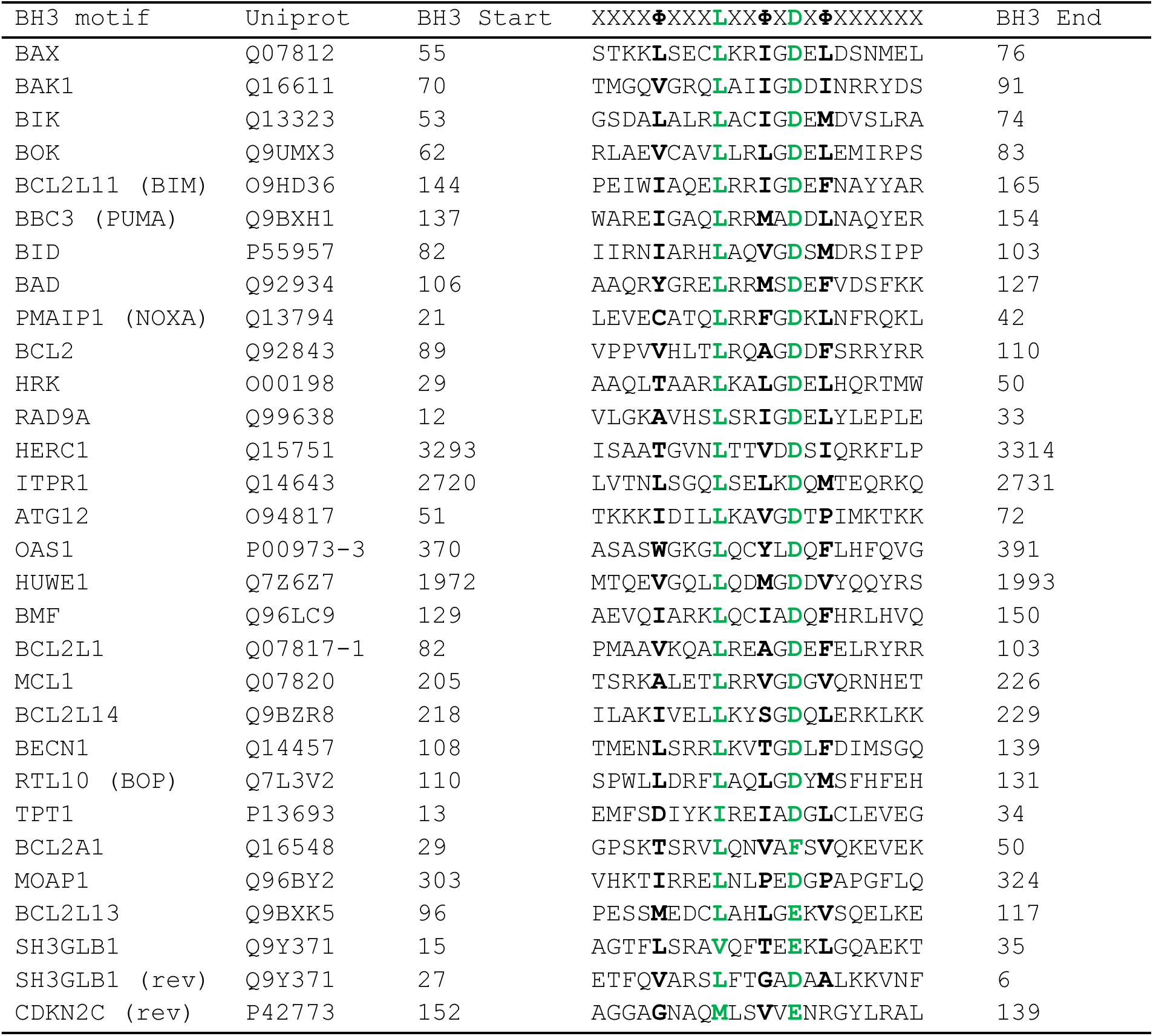
Here is an adapted version of the summary published by Sora and Papaleo in 2022 where the key conserved residues of each BH3 motif are colored in green and conserved hydrophobic residues are in bold. The final two peptides have reversed BH3 motifs.

## References

1. Ott, M., Robertson, J. D., Gogvadze, V., Zhivotovsky, B. & Orrenius, S. Cytochrome *c* release from mitochondria proceeds by a two-step process. Proceedings of the National Academy of Sciences 99, 1259–1263 (2002).

2. van Loo, G. et al. The serine protease Omi/HtrA2 is released from mitochondria during apoptosis. Omi interacts with caspase-inhibitor XIAP and induces enhanced caspase activity. Cell Death Differ. 9, 20–26 (2002).

3. Muñoz-Pinedo, C. et al. Different mitochondrial intermembrane space proteins are released during apoptosis in a manner that is coordinately initiated but can vary in duration. Proceedings of the National Academy of Sciences 103, 11573–11578 (2006).

4. Tait, S. W. G. & Green, D. R. Mitochondria and cell death: Outer membrane permeabilization and beyond. Nature Reviews Molecular Cell Biology vol. 11 621–632 Preprint at 10.1038/nrm2952 (2010).

5. Wolter, K. G. et al. Movement of Bax from the Cytosol to Mitochondria during Apoptosis. J. Cell Biol. 139, 1281–1292 (1997).

6. Hsu, Y.-T., Wolter, K. G. & Youle, R. J. Cytosol-to-membrane redistribution of Bax and Bcl-X L during apoptosis. Proceedings of the National Academy of Sciences 94, 3668–3672 (1997).

7. Desagher, S. et al. Bid-induced Conformational Change of Bax Is Responsible for Mitochondrial Cytochrome c Release during Apoptosis. J. Cell Biol. 144, 891–901 (1999).

8. Peyerl, F. W. et al. Elucidation of some Bax conformational changes through crystallization of an antibody–peptide complex. Cell Death Differ. 14, 447–452 (2007).

9. Nechushtan, A., Smith, C. L., Hsu, Y. & Youle, R. J. Conformation of the Bax C-terminus regulates subcellular location and cell death. EMBO J. 18, 2330–2341 (1999).

10. Gardai, S. J. et al. Phosphorylation of Bax Ser184 by Akt Regulates Its Activity and Apoptosis in Neutrophils. Journal of Biological Chemistry 279, 21085–21095 (2004).

11. Salvador-Gallego, R. et al. Bax assembly into rings and arcs in apoptotic mitochondria is linked to membrane pores. EMBO J. 35, 389–401 (2016).

12. Zhang, Y. et al. Structural basis of BAX pore formation. Science (1979). 388, (2025).

13. Wilfling, F. et al. BH3-only proteins are tail-anchored in the outer mitochondrial membrane and can initiate the activation of Bax. Cell Death Differ. 19, 1328–1336 (2012).

14. Czabotar, P. E. et al. Bax Crystal Structures Reveal How BH3 Domains Activate Bax and Nucleate Its Oligomerization to Induce Apoptosis. Cell 152, 519–531 (2013).

15. Schafer, B. et al. Mitochondrial Outer Membrane Proteins Assist Bid in Bax-mediated Lipidic Pore Formation. Mol. Biol. Cell 20, 2276–2285 (2009).

16. Montero, J. et al. Cholesterol and peroxidized cardiolipin in mitochondrial membrane properties, permeabilization and cell death. Biochimica et Biophysica Acta (BBA) - Bioenergetics 1797, 1217–1224 (2010).

17. Raemy, E. et al. Cardiolipin or MTCH2 can serve as tBID receptors during apoptosis. Cell Death Differ. 23, 1165–1174 (2016).

18. Dadsena, S. et al. Lipid unsaturation promotes BAX and BAK pore activity during apoptosis. Nat. Commun. 15, 4700 (2024).

19. Kuwana, T. et al. Bid, Bax, and Lipids Cooperate to Form Supramolecular Openings in the Outer Mitochondrial Membrane. Cell 111, 331–342 (2002).

20. Lutter, M. et al. Cardiolipin provides specificity for targeting of tBid to mitochondria. Nat. Cell Biol. 2, 754–756 (2000).

21. Lai, Y. C. et al. The role of cardiolipin in promoting the membrane pore-forming activity of BAX oligomers. Biochim. Biophys. Acta Biomembr. 1861, 268–280 (2019).

22. Li, Z. et al. The Mechanisms and Implications of Cardiolipin in the Regulation of Cell Death. Cell Biochem. Funct. 43, (2025).

23. Gallop, J. L. et al. Mechanism of endophilin N-BAR domain-mediated membrane curvature. EMBO J. 25, 2898–2910 (2006).

24. Bhatt, V. S., Ashley, R. & Sundborger-Lunna, A. Amphipathic Motifs Regulate N-BAR Protein Endophilin B1 Auto-inhibition and Drive Membrane Remodeling. Structure 29, 61–69.e3 (2021).

25. Karbowski, M., Jeong, S. Y. & Youle, R. J. Endophilin B1 is required for the maintenance of mitochondrial morphology. Journal of Cell Biology 166, 1027–1039 (2004).

26. Cuddeback, S. M. et al. Molecular cloning and characterization of Bif-1. A novel Src homology 3 domain-containing protein that associates with Bax. Journal of Biological Chemistry 276, 20559–20565 (2001).

27. Pierrat, B. et al. SH3GLB, a New Endophilin-Related Protein Family Featuring an SH3 Domain. Genomics 71, 222–234 (2001).

28. Takahashi, Y. et al. Loss of Bif-1 Suppresses Bax/Bak Conformational Change and Mitochondrial Apoptosis. Mol. Cell. Biol. 25, 9369–9382 (2005).

29. Etxebarria, A. et al. Endophilin B1/Bif-1 stimulates BAX activation independently from its capacity to produce large scale membrane morphological rearrangements. Journal of Biological Chemistry 284, 4200–4212 (2009).

30. Thorlacius, A., Rulev, M., Sundberg, O. & Sundborger-Lunna, A. Peripheral membrane protein endophilin B1 probes, perturbs and permeabilizes lipid bilayers. *Commun*. Biol. 8, 182 (2025).

31. Orlikowska-Rzeznik, H., Krok, E., Chattopadhyay, M., Lester, A. & Piatkowski, L. Laurdan Discerns Lipid Membrane Hydration and Cholesterol Content. J. Phys. Chem. B 127, 3382–3391 (2023).

32. Kagan, V. E. et al. Cytochrome c acts as a cardiolipin oxygenase required for release of proapoptotic factors. Nat. Chem. Biol. 1, 223–232 (2005).

33. Schug, Z. T. & Gottlieb, E. Cardiolipin acts as a mitochondrial signalling platform to launch apoptosis. Biochimica et Biophysica Acta (BBA) - Biomembranes 1788, 2022–2031 (2009).

34. Macdonald, P. J. et al. A dimeric equilibrium intermediate nucleates Drp1 reassembly on mitochondrial membranes for fission. Mol. Biol. Cell 25, 1905–1915 (2014).

35. Blaszczyk, M. et al. Modeling of protein–peptide interactions using the CABS-dock web server for binding site search and flexible docking. Methods 93, 72–83 (2016).

36. Abramson, J. et al. Accurate structure prediction of biomolecular interactions with AlphaFold 3. Nature 630, 493–500 (2024).

37. Große, L. et al. Bax assembles into large ring-like structures remodeling the mitochondrial outer membrane in apoptosis. EMBO J. 35, 402–413 (2016).

38. Schweighofer, S. V. et al. Endogenous BAX and BAK form mosaic rings of variable size and composition on apoptotic mitochondria. Cell Death Differ. 31, 469–478 (2024).

39. Taghiyev, A., Sun, D., Gao, Z. M., Liang, R. & Wang, L. Embelin-induced apoptosis of HepG2 human hepatocellular carcinoma cells and blockade of HepG2 cells in the G2/M phase via the mitochondrial pathway. Exp. Ther. Med. 4, 649–654 (2012).

40. Li, Y. et al. Embelin-induced MCF-7 breast cancer cell apoptosis and blockade of MCF-7 cells in the G2/M phase via the mitochondrial pathway. Oncol. Lett. 5, 1005–1009 (2013).

41. Park, N., Baek, H. S. & Chun, Y.-J. Embelin-Induced Apoptosis of Human Prostate Cancer Cells Is Mediated through Modulation of Akt and β-Catenin Signaling. PLoS One 10, e0134760 (2015).

42. Rostovtseva, T. K. et al. Bax activates endophilin B1 oligomerization and lipid membrane vesiculation. Journal of Biological Chemistry 284, 34390–34399 (2009).

43. Vannier, C., Pesty, A., San-Roman, M. J. & Schmidt, A. A. The Bin/Amphiphysin/Rvs (BAR) Domain Protein Endophilin B2 Interacts with Plectin and Controls Perinuclear Cytoskeletal Architecture. Journal of Biological Chemistry 288, 27619–27637 (2013).

44. Wang, Y. H. et al. Endophilin B2 promotes inner mitochondrial membrane degradation by forming heterodimers with Endophilin B1 during mitophagy. Sci. Rep. 6, (2016).

45. Serfass, J. M. et al. Endophilin B2 facilitates endosome maturation in response to growth factor stimulation, autophagy induction, and influenza A virus infection. Journal of Biological Chemistry 292, 10097–10111 (2017).

46. Yoshida, T. et al. Compartmentalization of soluble endocytic proteins in synaptic vesicle clusters by phase separation. iScience 26, 106826 (2023).

47. Mondal, S. et al. Multivalent interactions between molecular components involved in fast endophilin mediated endocytosis drive protein phase separation. Nature Communications 2022 13:1 13, 1–20 (2022).

48. Bademosi, A. T. et al. EndophilinA-dependent coupling between activity-induced calcium influx and synaptic autophagy is disrupted by a Parkinson-risk mutation. Neuron 111, 1402–1422.e13 (2023).

49. Decet, M. & Soukup, S.-F. Endophilin-A/SH3GL2 calcium switch for synaptic autophagy induction is impaired by a Parkinson’s risk variant. Autophagy 20, 925–927 (2024).

50. Hwang, M.-S. et al. Mitochondrial Ca2+ influx targets cardiolipin to disintegrate respiratory chain complex II for cell death induction. Cell Death Differ. 21, 1733–1745 (2014).

51. Schneider, C. A., Rasband, W. S. & Eliceiri, K. W. NIH Image to ImageJ: 25 years of image analysis. Nat. Methods 9, 671–675 (2012).

52. Punjani, A., Rubinstein, J. L., Fleet, D. J. & Brubaker, M. A. cryoSPARC: algorithms for rapid unsupervised cryo-EM structure determination. Nat. Methods 14, 290–296 (2017).

## Supplementary table reference

Sora, V. & Papaleo, E. (2022). Structural Details of BH3 Motifs and BH3-Mediated Interactions: An Updated Perspective. Front. Mol. Biosci., 9, 864874. 10.3389/fmolb.2022.864874

